# Meta-analysis reveals limited reproducibility of lncRNA-related findings in hepatocellular carcinoma research

**DOI:** 10.1101/2024.08.17.608405

**Authors:** Anamaria Necsulea, Philippe Veber, Tuyana Boldanova, Charlotte KY Ng, Stefan Wieland, Markus H Heim

## Abstract

**Background & Aims:** The human genome comprises tens of thousands of long non-coding RNAs (lncRNAs), whose functionality is highly debated. In the field of hepatocellular carcinoma (HCC) research, as for other cancer types, lncRNAs are increasingly reported to act as oncogenes or tumor suppressors and are put forward as useful diagnostic or prognostic biomarkers. Here, we investigate the reliability of these claims by performing a meta-analysis of the associations between HCC and lncRNAs reported in the scientific literature, and by assessing lncRNA expression patterns in two HCC patient cohorts.

**Approach & Results:** While nowadays up to 6% of all HCC-related publications cite lncRNAs, we show that most reported associations between HCC and lncRNAs have not been reproduced. In general, lncRNAs are less often differentially expressed between HCC tissues and controls compared to protein-coding genes. However, HCC-associated lncRNAs are frequently up-regulated in tumor samples, consistent with the fact that they are often selected based on transcriptome-wide comparative analyses. We perform a detailed examination of the 25 lncRNAs that are most frequently cited in association with HCC. For 10 out of these 25 lncRNAs, including well known lncRNAs such as *MALAT1*, *NEAT1*, *H19* and *XIST*, we identify important conflicts between the biological roles and expression patterns previously reported for them in HCC and the expression patterns that we observe here. Finally, we observe that HCC-associated publications that cite lncRNAs are retracted three times more often than publications that cite protein-coding genes.

**Conclusions:** Our results thus highlight the poor reproducibility of lncRNA-related claims in association with HCC, which is problematic in a context where new biomarkers and molecular targets for therapy are greatly needed.

## Introduction

The human genome harbors tens of thousands of long non-coding RNA (lncRNA) genes (Amaral et al. 2023). These transcripts are simply defined as long RNA molecules (at least 200 nucleotides) that do not encode functional proteins. Long non-coding RNAs with major biological functions have been known for several decades (Brannan et al. 1990; Brown et al. 1991), well before the magnitude of the lncRNA gene repertoire was first perceived (Guttman et al. 2009; Djebali et al. 2012; Derrien et al. 2012). Numerous recent studies have proposed that lncRNAs play important roles in gene expression regulation, genome stability or nuclear architecture (Engreitz, Ollikainen, and Guttman 2016). Because of these promising findings, a great deal of effort has been put into investigating the contributions of lncRNAs to cancer biology (Gutschner and Diederichs 2012). The interest in lncRNAs is even stronger for cancer types for which effective drug targets and disease biomarkers are still urgently needed, such as hepatocellular carcinoma (HCC).

Hepatocellular carcinoma (HCC) is one of the most frequent causes of cancer-related mortality (J. D. Yang et al. 2019). As HCC is generally detected at late stages of tumor progression, surgical treatment options are unavailable for the majority of patients (Hartke, Johnson, and Ghabril 2017). Several systemic therapies now exist, but they increase median patient survival by less than 1 year (Finn et al. 2020). Thus, developing new treatments and biomarkers for the early detection of HCC is imperative. With this goal, there has been extensive research aiming to identify the genomic, transcriptomic and proteomic features that are altered in HCC (The Cancer Genome Atlas Research Network 2017; Jiang et al. 2019; Pinyol et al. 2021; Ng et al. 2022). In particular, hundreds of differentially expressed lncRNAs were identified by large-scale transcriptomics studies that compared HCC samples with adjacent non-tumor tissue or with normal liver samples (Cui et al. 2017; Y. Yang et al. 2017; Li et al. 2019; Juan P. Unfried et al. 2019). Some of the lncRNAs associated with HCC were subject to further experimental investigations, aiming to elucidate their mechanisms of action and the consequences of their differential regulation in tumors. As a result, many lncRNAs were proposed to act as oncogenes or as tumor suppressors in hepatocellular carcinoma (Lanzafame et al. 2018; Juan Pablo Unfried et al. 2021). For some lncRNAs, experimental analyses of their roles in cancer have led to conflicting results. For example, the *H19* lncRNA was alternatively proposed to act as a tumor suppressor (Hao et al. 1993; Yoshimizu et al. 2008; Schultheiss et al. 2017) or as an oncogene (Matouk et al. 2007; Zhou et al. 2019) in various cancer types including HCC (Tietze and Kessler 2020). Likewise, *MALAT1*, initially described as an abundant lncRNA associated with the presence of metastases (Ji et al. 2003), was first proposed to promote tumor growth and invasion in breast cancer (Arun et al. 2016), but is now believed to be a tumor suppressor (Kim et al. 2018).

While conflict and resolution are part of the normal scientific process, contradictions regarding the roles of lncRNAs in HCC and other cancer types need to be interpreted in light of the existing debates in the lncRNA research field. The functionality of lncRNAs has been contested since the discovery that they are overall poorly conserved during evolution (Haerty and Ponting 2014). As an example, only about 10% of human lncRNAs are shared with mouse (Necsulea et al. 2014; Sarropoulos et al. 2019). The low rate of lncRNA evolutionary conservation supports the hypothesis that the majority of lncRNAs are non-functional. The opposite view is that lncRNAs can be biologically functional in the absence of evolutionary conservation (Mattick et al. 2023), in contradiction with evolutionary theory (Haerty and Ponting 2014; Ponting and Haerty 2022). Beyond the issue of evolutionary conservation, functional validations of lncRNAs in the lab have also yielded controversial results. Ascertaining lncRNA functions is challenging, in part because genomic *loci* that give rise to lncRNAs can also contain additional functional genomic elements that are independent of the transcription of the lncRNA (Bassett et al. 2014). Genetic perturbations of lncRNA *loci* can thus have phenotypic consequences that are not due to the absence of the RNA molecule. In agreement with this, in several noteworthy cases, biological functions that were originally attributed to the lncRNA molecule following genetic alterations of lncRNA *loci* were subsequently proven to stem from the presence of DNA sequences involved in expression regulation at the *locus* (Groff et al. 2016; 2018). In other cases, there were contradictions between lncRNA functional assays performed *in vivo* and *in vitro*, or with knockout and knockdown approaches (Goudarzi et al. 2019; Amândio et al. 2016; Kim et al. 2018). These inconsistencies can sometimes stem from the lack of appropriate controls in lncRNA functional experiments (Kim et al. 2018).

The controversies related to lncRNA functions are likely to become resolved with time, as the research field becomes more mature. However, cancer research cannot afford to wait to assess the validity of the reported roles of lncRNAs. Here, we addressed this issue by evaluating the reproducibility of lncRNA-related claims in hepatocellular carcinoma. We scanned the scientific literature to extract lncRNAs that have been associated with HCC and we examined their characteristics, in comparison with protein-coding genes and with other lncRNAs. We assessed their expression patterns in two large HCC patient cohorts. We examined in detail the 25 lncRNAs with the highest numbers of HCC-associated citations and we highlight several contradictions between the roles that were previously attributed to these lncRNAs and the expression patterns that we observe here.

## Results

### lncRNAs are often associated with HCC in the scientific literature

We first aimed to evaluate the prevalence of lncRNA studies in the HCC-related scientific literature. To do this, we performed a PubMed search for scientific articles that have the term “hepatocellular carcinoma” in the title (Supplementary Table 1). As lncRNAs were only detected at the genome-wide level thanks to the advent of RNA sequencing, we restricted our analysis to articles published between 2009 and 2023, obtaining a total of 46,960 publications. We extracted the gene names that were cited in the abstracts of the articles and matched them with HGNC gene symbols for protein-coding genes and lncRNAs, as listed in the Ensembl database (Materials and methods). Although we cannot manually verify the roles attributed to these genes in all publications, the fact that these genes were cited in the abstracts indicates that they were deemed to be important for the main messages of the articles. We observed that the frequency of lncRNA-citing HCC articles increased rapidly between 2009 and 2020, starting from 0.1% in 2009 and reaching a maximum of 5.6% in 2020 (Figure 1a). The frequency decreased in 2021 and over the next two years, reaching 2.16% in 2023. In total, 574 lncRNAs were cited in the abstracts of HCC-related publications. We found that the majority (54%) of these lncRNAs were cited in only one publication, suggesting that most lncRNA-related claims in HCC have not yet been reproduced (Figure 1b, Supplementary Table 2). Only 33 lncRNAs were cited in 10 or more HCC-related publications. In contrast, the frequency of HCC-related articles that cite protein-coding genes is more stable with time, varying only between 44% and 57% in the same period (Supplementary Figure 1a). In total, 7,163 protein-coding genes are cited in HCC-related articles, the majority of which (60.7%) are cited in 2 or more publications (Supplementary Figure 1b, Supplementary Table 3). These results highlight the contrasting dynamics of the HCC scientific literature with respect to protein-coding genes and lncRNAs.

**Figure 1.**
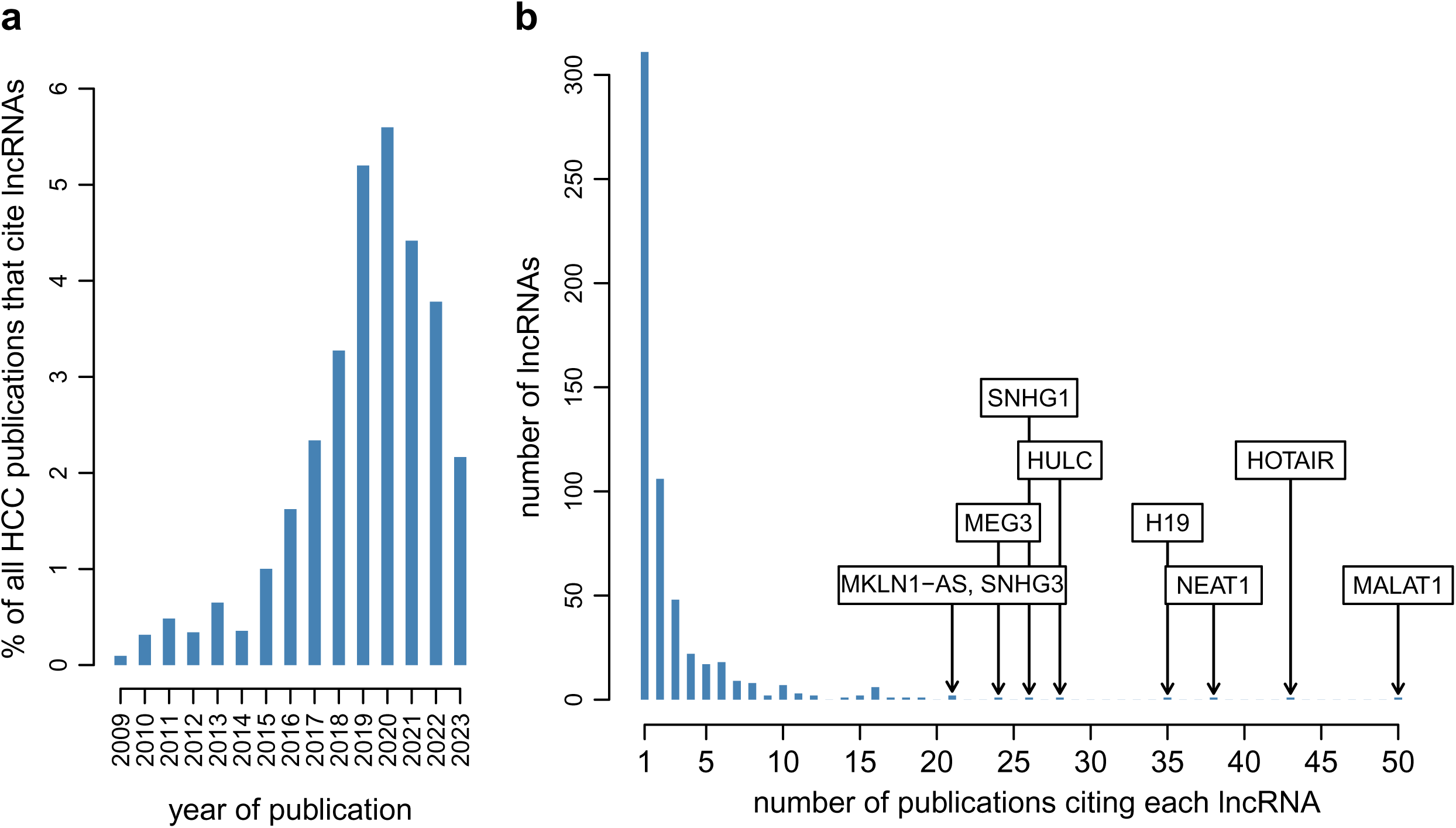
Long non-coding RNAs are frequently cited in association with HCC. a) Barplot representing the percentage of all HCC publications that cite long non-coding RNAs in the abstract, between 2009 and 2022. b) Histogram of the number of HCC-associated publications in which each lncRNA is cited in the abstract. Labels indicate the names of the lncRNAs with 20 or more citations.

### Characteristics of HCC-associated lncRNAs

We next evaluated the genomic and transcriptomic characteristics of the lncRNAs that were associated with HCC in scientific publications. We divided protein-coding genes and lncRNAs into three classes: those that are cited in more than 1 HCC-associated articles (below, we refer to these genes as “frequently cited”), those that are cited in exactly 1 HCC-associated article (below termed “infrequently cited”), and those that are not cited in association with HCC (below termed “not cited” or “other”, Materials and methods). We first analyzed the expression levels for the 6 categories of genes in HCC tumors and adjacent tissues, using transcriptome sequencing data from two large HCC patient cohorts (Ng et al. 2022; The Cancer Genome Atlas Research Network 2017). The first transcriptome collection comprises RNA-seq data derived from paired tumor and adjacent tissue biopsies, for 105 HCC patients for which extensive clinical information was compiled (Ng et al. 2022). Sample characteristics are provided in Supplementary Tables 4 and 5 for tumor and adjacent tissue, respectively. The second dataset corresponds to the widely used TCGA resource (The Cancer Genome Atlas Research Network 2017). We focused on cases for which paired tumor and adjacent tissue samples were available and could thus analyze data from 37 patients in TCGA. We computed the average expression levels across tumor samples and across adjacent tissue samples, for each gene and for each patient cohort. Irrespective of the number of HCC-associated citations, average expression levels were significantly lower for lncRNAs than for protein-coding genes, for both tissue types and for both patient cohorts (Figure 2a,b, Supplementary Figure 2, Wilcoxon rank sum test, p-value < 1e-10 for all comparisons). Likewise, we observed that frequently cited lncRNAs had significantly higher expression levels than infrequently cited lncRNAs, which in turn had significantly higher expression levels than those that were not cited (Wilcoxon rank sum test, p-value < 1e-10 for all comparisons except for lncRNAs cited once *vs.* lncRNAs cited at least 2 times in adjacent tissue samples, p-value 4e-9). The same conclusions were reached for the two transcriptome datasets (Figure 2, Supplementary Figure 2).

**Figure 2.**
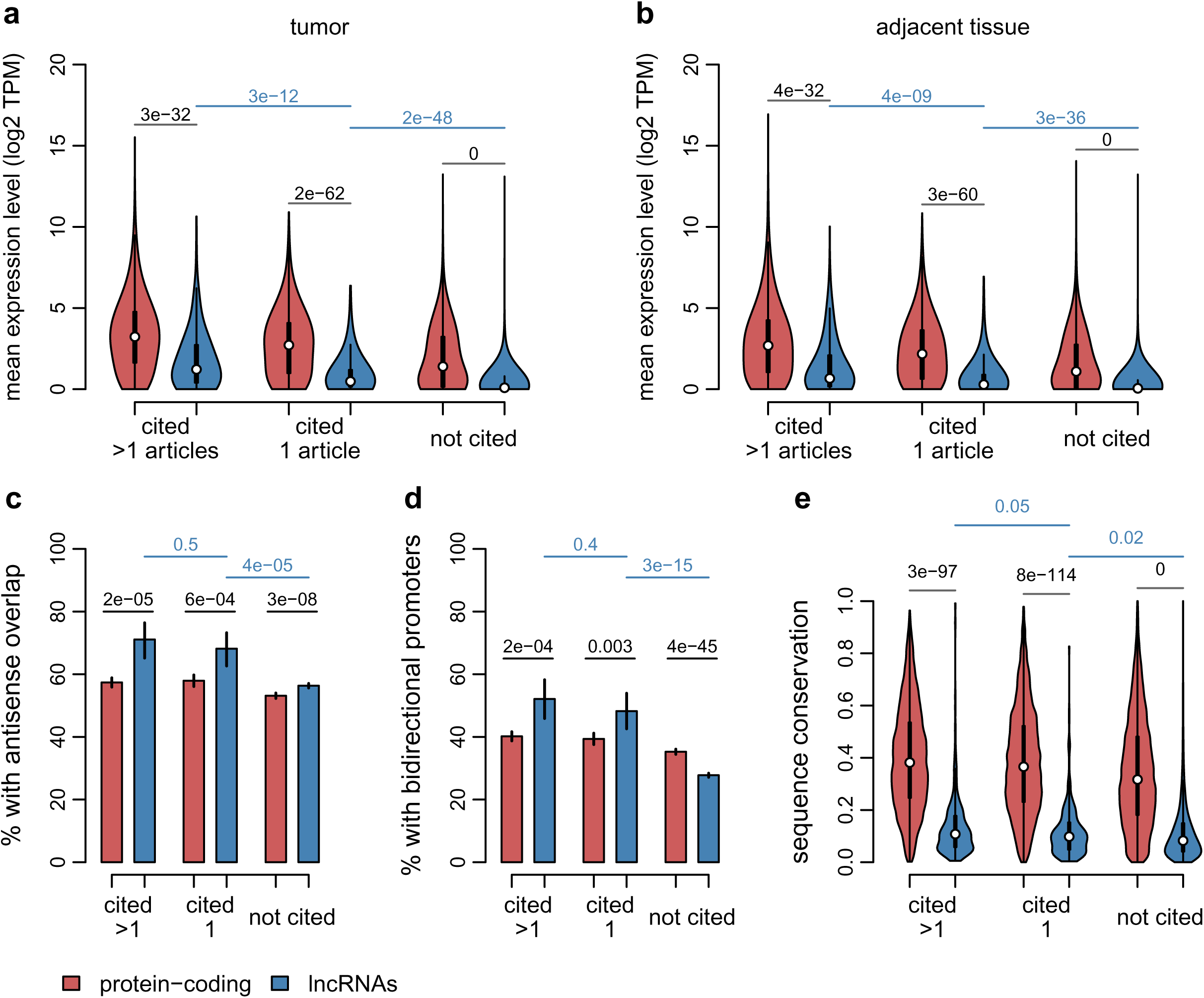
Characteristics of HCC-associated lncRNAs and comparison with other lncRNAs and with protein-coding genes. Protein-coding genes are depicted in red and divided into 3 categories: cited in more than one HCC publication, in exactly one publication, or not cited. Long non-coding RNAs are depicted in blue and divided in the same categories. Red: protein-coding genes. Blue: lncRNAs. a) Violin plots representing the distribution of the average expression level (log2-transformed TPM) for Ng et al 2022 tumor samples. b) Same as a), for Ng et al 2022 adjacent tissue samples. c) Percentage of genes that have antisense overlap with other genes. d) Percentage of genes that have a bidirectional promoter. e) Average exonic sequence conservation score (PhastCons score). a) to e) Numeric values represent p-values of the Wilcoxon rank sum test (panels a, b and e) or of the Chi-squared test (panels c, d), for the comparisons between protein-coding genes and lncRNAs, or between citation classes for lncRNAs. For the barplots, vertical bars represent 95% confidence intervals.

We also observed that lncRNAs that are cited in association with HCC have particular genomic characteristics. They overlap more frequently with other genes on the antisense strand than other lncRNAs (71.1% for frequently cited lncRNAs, 68% for infrequently cited lncRNAs and 56% for lncRNAs that are not cited, Figure 2c). They are also more often transcribed from bidirectional promoters than other lncRNAs (52% for lncRNAs cited more than once, 48% for lncRNAs cited exactly once and 28% for lncRNAs not cited, Figure 2d). The differences are statistically significant for the comparison between lncRNAs that are cited at least once and lncRNAs that are not cited in HCC (Chi-squared test, p-value 4e-5 for the antisense overlaps, p-value < 1e-10 for the presence of bidirectional promoters, Figure 2c,d). Importantly, the proportions of *loci* with antisense overlaps and with bidirectional promoters are also significantly higher for lncRNAs than for protein-coding genes for the two first citation classes (Figure 2c,d). This pattern is less striking or not observed for genes that are not cited in association with HCC. In particular, in this last subcategory, there is a tendency in the other direction: lncRNAs have significantly fewer bidirectional promoters than protein-coding genes (Chi-squared test, p-value < 1e-10, Figure 2d). Thus, lncRNAs that are associated with HCC in scientific publications have different genomic characteristics than other lncRNAs and than protein-coding genes (Supplementary Tables 6 and 7).

Finally, we evaluated the extent of exonic sequence conservation for the different categories of genes. We used as a basis the PhastCons score (Siepel et al. 2005), computed on a whole genome alignment between human and 29 other mammals (Materials and methods). Given the frequent overlaps between lncRNAs and other genes, we used for this analysis only those exonic regions that do not overlap with exons from other genes. We observed that, irrespective of their citation status in association with HCC, exonic sequence conservation levels are overwhelmingly higher for protein-coding genes than for lncRNAs (Wilcoxon rank sum test, p-value < 1e-10 for all three comparisons, Figure 2e). For protein-coding genes, sequence conservation scores were significantly higher for genes that were cited in association with HCC than for genes that were not cited (median score 0.38 for frequently cited genes, 0.37 for infrequently cited genes and 0.32 for other genes, Wilcoxon rank sum test, p-value < 1e-10 for the comparison between the last two categories). For lncRNAs, the difference between citation classes was only marginally significant (median score 0.11 for frequently cited lncRNAs, 0.10 for infrequently cited lncRNAs and 0.08 for other lncRNAs, Wilcoxon rank sum test, p-value 0.024 for the comparison between the last two categories, Figure 2e).

### HCC-associated lncRNA expression patterns

We then analyzed gene expression patterns in HCC tissues, for lncRNAs and protein-coding genes. We first performed principal component analyses (PCA) based on the expression levels of protein-coding genes and lncRNAs, divided into citation classes as above, for the two patient cohorts. Samples were grouped according to tissue types and to tumor differentiation grades on the first axis of all PCAs (Supplementary Figures 3 and 4). This shows that expression patterns of both gene categories were sufficient to distinguish between tumor and adjacent tissues and between tumor differentiation grades, irrespective of their association with HCC in the literature. However, the amount of variance explained by the first PCA axis was generally higher for protein-coding genes than for lncRNAs, and for frequently cited lncRNAs than for other lncRNAs (Supplementary Figures 3 and 4), indicating a higher discriminative power for these gene categories.

We next performed two differential expression (DE) analyses. First, we compared expression levels between paired tumor and adjacent tissue samples, for the two patient cohorts (Ng et al. 2022; The Cancer Genome Atlas Research Network 2017). Second, we compared expression levels between tumor samples depending on their degree of differentiation, as evaluated by the Edmondson-Steiner grade (Martins-Filho et al. 2017), in the first patient cohort (Ng et al. 2022). We identified significantly differentially expressed genes by setting a maximum false discovery rate (FDR) threshold of 5% (Supplementary Tables 8 to 10). We compared the frequency of significantly differentially expressed genes between protein-coding genes and lncRNAs, within each citation class. We observed different tendencies for upregulated and downregulated genes. First, we found that frequently cited lncRNAs are upregulated in tumors compared to adjacent tissues at least as often as protein-coding genes, within the same citation class (Figure 3a). For the TCGA cohort, frequently cited lncRNAs are significantly more often upregulated in tumors compared to adjacent tissues than protein-coding genes (Chi-squared test, p-value 0.002, Supplementary Figure 5). Likewise, frequently cited lncRNAs are upregulated in tumors with high grades compared to tumors with low grades as often as protein-coding genes (Figure 3c). These patterns are less strong for infrequently cited lncRNAs, and not observed for lncRNAs that are not cited, which are significantly less often upregulated than protein-coding genes, for both DE analyses (Chi-squared test, p-value < 1e-10, Figure 3a,c). Second, in contrast with the observations for upregulated genes, we observed that lncRNAs are less often downregulated than protein-coding genes, for all citation classes and for both DE analyses (Chi-squared test, p-value < 0.02 for all comparisons, Figure 3b,d). Finally, we observed that for both DE analyses and for both DE patterns (upregulation and downregulation), lncRNAs that are cited in association with HCC are more often significantly differentially expressed than lncRNAs that are not cited (Chi-squared test, p-value 0.004 for downregulation in tumors with high grades, p-value < 1e-10 in all other cases, Figure 3, Supplementary Figure 5). Overall, these observations are expected, given that lncRNAs are often associated with HCC in publications because they are identified as differentially expressed between HCC tissues and controls in transcriptomic data analyses. We also analyzed the consistency of DE patterns between the two cohorts, for the same categories of genes (Supplementary Figure 6). We observed that DE patterns are more often consistent between cohorts for protein-coding genes than for lncRNAs, irrespective of their citation class (Supplementary Figure 6).

**Figure 3.**
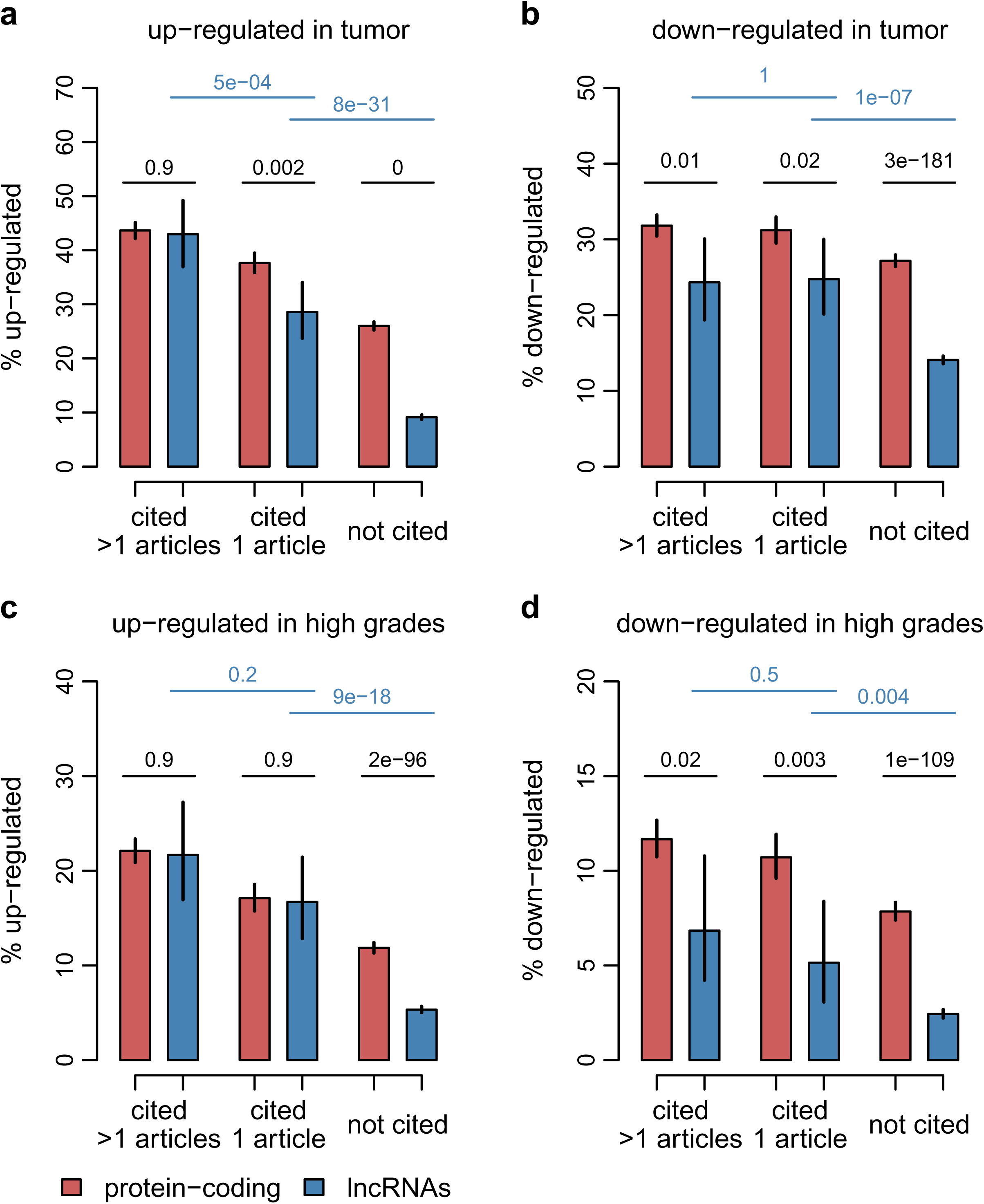
Differential expression patterns for HCC-associated lncRNAs and comparison with other lncRNAs and with protein-coding genes, in the Ng et al 2022 transcriptome dataset. Protein-coding genes are depicted in red and divided into 3 categories: cited in more than one HCC publication, in exactly one publication, or not cited. Long non-coding RNAs are depicted in blue and divided in the same categories. A maximum FDR threshold was set to extract significantly DE genes. a) Percentage of genes that are significantly up-regulated in tumors compared to adjacent tissues. b) Percentage of genes that are significantly down-regulated in tumors compared to adjacent tissues. c) Percentage of genes that are significantly up-regulated in tumor samples with Edmonson-Steiner grades III and IV compared with tumor samples with Edmondson-Steiner grades I and II. e) Percentage of genes that are significantly down-regulated in tumor samples with Edmonson-Steiner grades III and IV compared with tumor samples with Edmondson-Steiner grades I and II. a) to e) Numeric values represent p-values of the Chi-squared test for the comparisons between protein-coding genes and lncRNAs, or between citation classes for lncRNAs. Vertical bars represent 95% confidence intervals.

Having shown that HCC-associated lncRNAs are more often transcribed from bidirectional promoters than protein-coding genes, we wanted to test whether in these cases the neighboring genes also had altered expression in HCC. Indeed, we found that up to 30% of HCC-associated lncRNAs (frequently or infrequently cited) had a close neighbor that was transcribed from the same bidirectional promoter and that was significantly differentially expressed between tumors and adjacent issues, for both patient cohorts (Supplementary Figure 7). These proportions were significantly higher than for HCC-associated protein-coding genes (Chi-squared test, p-value < 1e-10), and for lncRNAs that are not associated with HCC (Chi-squared test, p-value 5e-7 for the TCGA analysis, p-value < 1e-10 in all other cases, Supplementary Figure 7). The same conclusion was reached for the DE analysis comparing tumor differentiation grades (Supplementary Figure 7). These results suggest that lncRNAs found in close vicinity to DE protein-coding genes are more often investigated in HCC publications.

### Inconsistent expression patterns for lncRNAs that are frequently associated with HCC

We next wanted to investigate in detail the expression patterns of lncRNAs that are the most frequently cited in association with HCC. We selected the 25 lncRNAs that had the highest number of HCC-associated citations. These include *MALAT1*, *HOTAIR*, *NEAT1*, *H19*, *XIST* and other lncRNAs that are often discussed in cancer research (Figure 4, Supplementary Figure 8, Table 1). For these 25 lncRNAs, we browsed the literature to extract their previously reported biological roles and expression patterns in HCC (Materials and methods, Table 1, Supplementary Table 11). We compared these previously reported observations with our own findings regarding the differential expression of these lncRNAs in the two patient cohorts (Table 1). We found inconsistencies for several of the most prominent lncRNAs.

**Figure 4.**
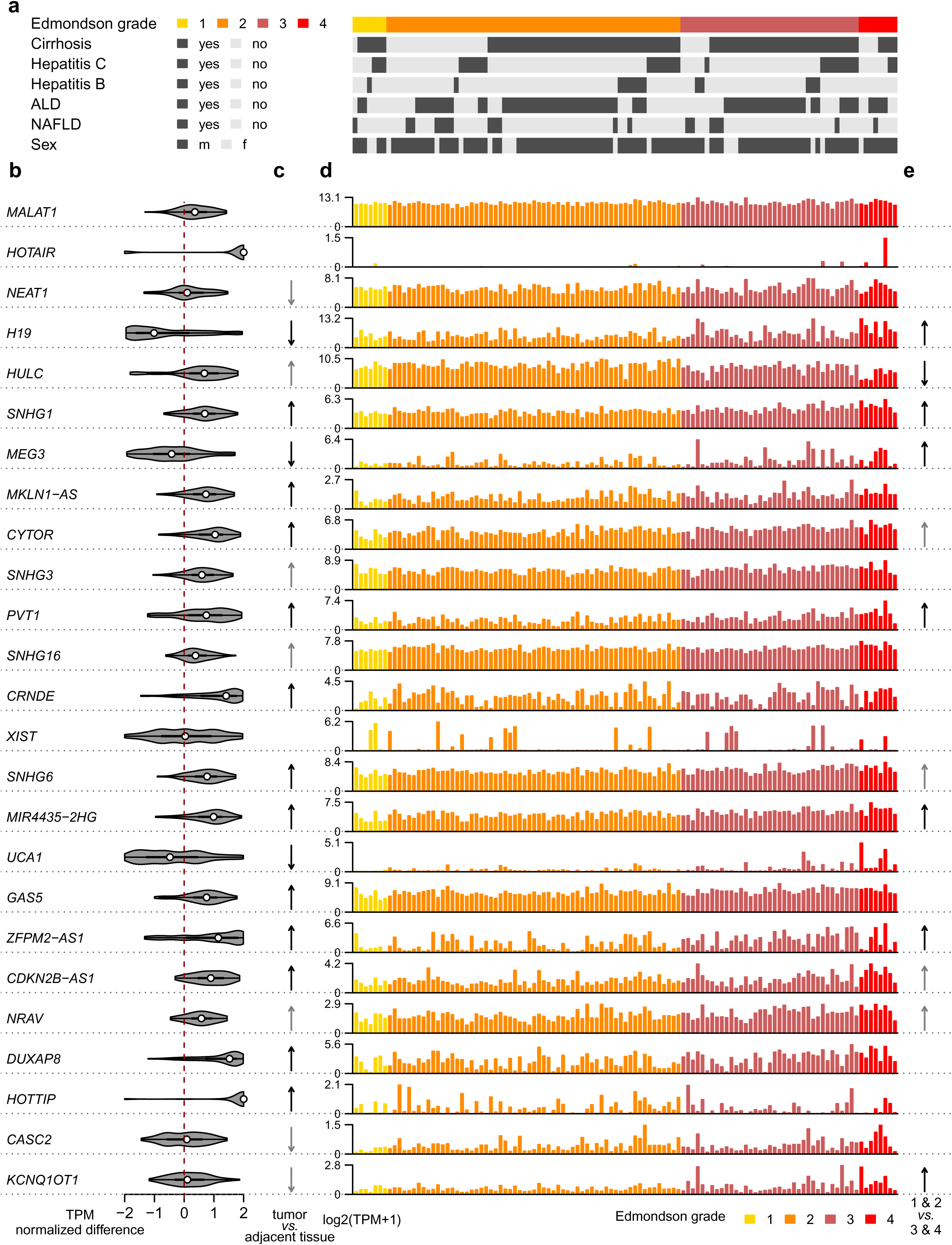
Expression patterns for the 25 lncRNAs with the highest numbers of HCC-associated citations, in the Ng et al 2022 transcriptome dataset. a) Clinical characteristics for the tumor samples included in this analysis. b) Violin plot depicting the distribution of the normalized expression differences between paired tumor and adjacent tissue biopsies, across the 105 analyzed patients. c) Arrows indicating whether the difference between tumor and paired adjacent tissue biopsies was statistically significant (FDR < 5%), and if so, the direction of the expression change (up-regulated or down-regulated). Dark gray arrows indicate absolute fold expression changes above 1.5. d) Barplot depicting the expression levels (log2-transformed TPM) across the tumor samples. Bars are colored depending on the Edmondson-Steiner grade of the tumors. e) Arrows indicating whether the difference between tumor and paired adjacent tissue biopsies was statistically significant (FDR < 5%), and if so, the direction of the expression change (up-regulated or down-regulated).

**Table 1.** Characteristics of the 25 lncRNAs that are most frequently cited in association with HCC. The lncRNAs for which inconsistencies were detected are highlighted in bold. a) Expression pattern in the Ng et al 2022 cohort, in the DE analysis contrasting tumor biopsies with adjacent tissue biopsies. b) Expression pattern in the TCGA HCC cohort, in the DE analysis contrasting tumor biopsies with adjacent tissue biopsies. c) Number of articles focusing on HCC and lncRNAs that cite the focus lncRNA in the abstract, excluding reviews. d) Expression patterns reported in the literature, for the comparison between HCC tumor tissues and adjacent tissue or normal liver. Studies performed exclusively on cell lines are excluded. e) Reported role for the focal lncRNA (oncogene or tumor suppressor) in the literature. f) Number of reported regulatory interactions between the focal lncRNA and microRNAs, as a competing endogenous RNA (ceRNA or miRNA sponge). a,b) A maximum FDR threshold of 5% was set to identify significantly differentially expressed genes.

| Name | Ng et al 2022 expression <sup>a</sup> | TCGA expression <sup>b</sup> | N <sup>c</sup> | Reported expression <sup>d</sup> | Reported role <sup>e</sup> | Reported ceRNA interactions <sup>f</sup> |
| --- | --- | --- | --- | --- | --- | --- |
| *MALAT1* | N.S. | upregulated | 32 | upregulated (n=14)<br>N.S. (n=3) | oncogene (n=12) | 10 |
| *HOTAIR* | N.S. | N.S. | 26 | upregulated (n=12) | oncogene (n=7) | 2 |
| *NEAT1* | downregulated | N.S. | 32 | upregulated (n=18) | oncogene (n=12) | 4 |
| *H19* | downregulated | downregulated | 20 | upregulated (n=3)<br>downregulated (n=3)<br>N.S. (n=1) | oncogene (n=6)<br>tumor suppressor (n=2) | 3 |
| *HULC* | upregulated | upregulated | 18 | upregulated (n=6) | oncogene (n=3) | 6 |
| *SNHG1* | upregulated | upregulated | 20 | upregulated (n=9) | oncogene (n=4) | 10 |
| *MEG3* | downregulated | downregulated | 17 | downregulated (n=7) | oncogene (n=1)<br>tumor suppressor (n=2) | 4 |
| *MKLN1-AS* | upregulated | upregulated | 19 | upregulated (n=6) | oncogene (n=1) | 2 |
| *CYTOR* | upregulated | upregulated | 18 | upregulated (n=7) | oncogene (n=6) | 6 |
| *SNHG3* | upregulated | upregulated | 15 | upregulated (n=4) | oncogene (n=3) | 1 |
| *PVT1* | upregulated | upregulated | 15 | upregulated (n=9) | oncogene (n=1) | 3 |
| *SNHG16* | upregulated | N.S. | 14 | upregulated (n=11) | oncogene (n=2) | 7 |
| *CRNDE* | upregulated | upregulated | 16 | upregulated (n=11) | oncogene (n=5) | 3 |
| *XIST* | N.S. | N.S. | 13 | upregulated (n=5)<br>downregulated (n=4) | oncogene (n=4)<br>tumor suppressor (n=2) | 4 |
| *SNHG6* | upregulated | upregulated | 11 | upregulated (n=4) | oncogene (n=5) | 4 |
| *MIR4435-2HG* | upregulated | upregulated | 12 | upregulated (n=7) | oncogene (n=5) | 3 |
| *UCA1* | downregulated | downregulated | 8 | upregulated (n=3) | oncogene (n=3) | 2 |
| *GAS5* | upregulated | upregulated | 10 | upregulated (n=2)<br>downregulated (n=4) | oncogene (n=1)<br>tumor suppressor (n=3) | 4 |
| *ZFPM2-AS1* | upregulated | upregulated | 14 | upregulated (n=5) | oncogene (n=4) | 2 |
| *CDKN2B-AS1* | upregulated | upregulated | 12 | upregulated (n=8) | oncogene (n=5) | 4 |
| *NRAV* | upregulated | upregulated | 13 | upregulated (n=2) | oncogene (n=1) | 1 |
| *DUXAP8* | upregulated | upregulated | 11 | upregulated (n=5) | oncogene (n=4) | 4 |
| *HOTTIP* | upregulated | upregulated | 8 | upregulated (n=3) | oncogene (n=4) | 0 |
| *CASC2* | downregulated | N.S. | 10 | downregulated (n=8) | tumor suppressor (n=5) | 3 |
| *KCNQ1OT1* | downregulated | N.S. | 8 | upregulated (n=5) | oncogene (n=5) | 5 |

First, there were contradictory claims in the literature regarding the expression patterns of *MALAT1*, *H19*, *XIST* and *GAS5* (Table 1, Figure 4, Supplementary Figure 8). *MALAT1* was reported as upregulated in HCC tumors compared to controls in 14 studies and not significantly differentially expressed in 3 studies. Our analyses show that it is upregulated in HCC tumors compared to controls in the TCGA cohort, but not in the other patient cohort that we analyzed. For *H19*, previous publications report it either as upregulated (3 studies), downregulated (3 studies) or not significantly DE (1 study) between HCC tumors and controls, while we find that it is significantly downregulated in tumors compared to adjacent tissue samples. *XIST* was reported to be upregulated in HCC tissues compared to controls in 4 studies and downregulated in 3 studies, while we find that it is not significantly differentially expressed between tissue types, for both patient cohorts. Importantly, we note that our DE analyses include the sex of the patient as a factor (Materials and methods), which is particularly relevant for *XIST* given its predominant expression in females. Finally, *GAS5* was reported as upregulated in HCC tumors compared to controls (2 studies) and downregulated (4 studies), while we found it to be upregulated in tumors compared to adjacent tissues, for both cohorts.

Second, we observed contradictions between previous reports and our own results for *HOTAIR*, *NEAT1*, *UCA1* and *KCNQ1OT1* (Table 1, Figure 4, Supplementary Figure 8). In the case of *HOTAIR*, all 12 previous studies report it as up-regulated in HCC tissues compared to controls, while we find that it is not significantly DE between tumors and adjacent tissues, nor between tumor differentiation grades. *NEAT1* and *KCNQ1OT1* were reported to be upregulated in HCC tumors compared to controls in all available studies (18 for *NEAT1* and 5 for *KCNQ1OT1)*, while we found them to be significantly downregulated in one patient cohort and not significantly DE in the other one. *UCA1* was reported as upregulated in the literature (3 studies), while we found it to be significantly downregulated in both patient cohorts. To a lesser extent, we observe inconsistencies for *SNHG16* and *CASC2*, for which we only found a significant DE pattern in one patient cohort (Table 1, Figure 4, Supplementary Figure 8).

In addition to the reported expression patterns, we also analyzed the biological roles attributed to lncRNAs in the literature. We observed that the top 25 lncRNAs were often claimed to act as oncogenes or tumor suppressors, depending on their expression pattern. Almost systematically, lncRNAs that were upregulated in tumors compared to controls were reported to act as oncogenes, while downregulated lncRNAs were reported to act as tumor suppressors. One exception is *MEG3*, for which an oncogene role was claimed in one publication, despite its downregulation in tumors (Table 1). We were not able to verify whether the oncogene or tumor suppressor roles were supported by additional experimental evidence. We also observed that almost all of the top 25 lncRNAs were reported to act as competing endogenous RNAs (Ebert and Sharp 2010), also known as sponges or decoy targets for microRNAs (Table 1). The only exception was *HOTTIP* (Table 1).

## Discussion

In this manuscript, our main goal was to investigate the reproducibility of lncRNA findings in HCC. We performed a literature search which allowed us to evaluate the frequency of articles that prominently cite lncRNAs in association with HCC (Materials and methods). This frequency has rapidly increased since 2009, reaching almost 6% of all HCC-related articles in 2020. This phenomenon is at least in part due to the fact that lncRNAs became easy to detect around 2009, thanks to the development of sensitive transcriptome sequencing technologies (Wang, Gerstein, and Snyder 2009). Interestingly, the frequency of lncRNA-citing articles among all HCC-associated publications strongly decreased between 2021 and 2023. The fact that the peak frequency was reached the year before the Covid-19 pandemic might perhaps be attributed to a shift in interest, moving away from lncRNAs and perhaps towards topics related to Covid-19. While this shift is understandable, this could suggest that the growing interest in lncRNAs in HCC research may have been to some extent due to a fashion effect, which could easily decline if other “hot topics” appear. The decline in the frequency of lncRNA-citing articles could also be due to the growing interest for immunotherapy in HCC (Sangro et al., 2021), as these new treatments do not target lncRNAs.

We found that the majority of HCC-associated lncRNAs were cited by exactly one article, as might be expected given that lncRNAs are still a fairly new research topic. Nevertheless, this observation is important, as it indicates that most lncRNA-related findings in HCC have not yet been reproduced and that they should be considered with great caution. With this in mind, we analyzed the frequency of retractions for lncRNA-citing articles (Figure 5, Materials and methods). We found that 3.25% of all lncRNA-citing articles were retracted, which is almost three times more than the frequency observed for articles that cite protein-coding genes (1.16%, Chi-squared test, p-value 5.5e-10). This pattern holds even when articles are analyzed by year of publication (Figure 5b). Given the time lag between an initial publication and a potential retraction, we predict that the number of retractions of lncRNA-citing articles in HCC will increase even more in the next few years.

**Figure 5.**
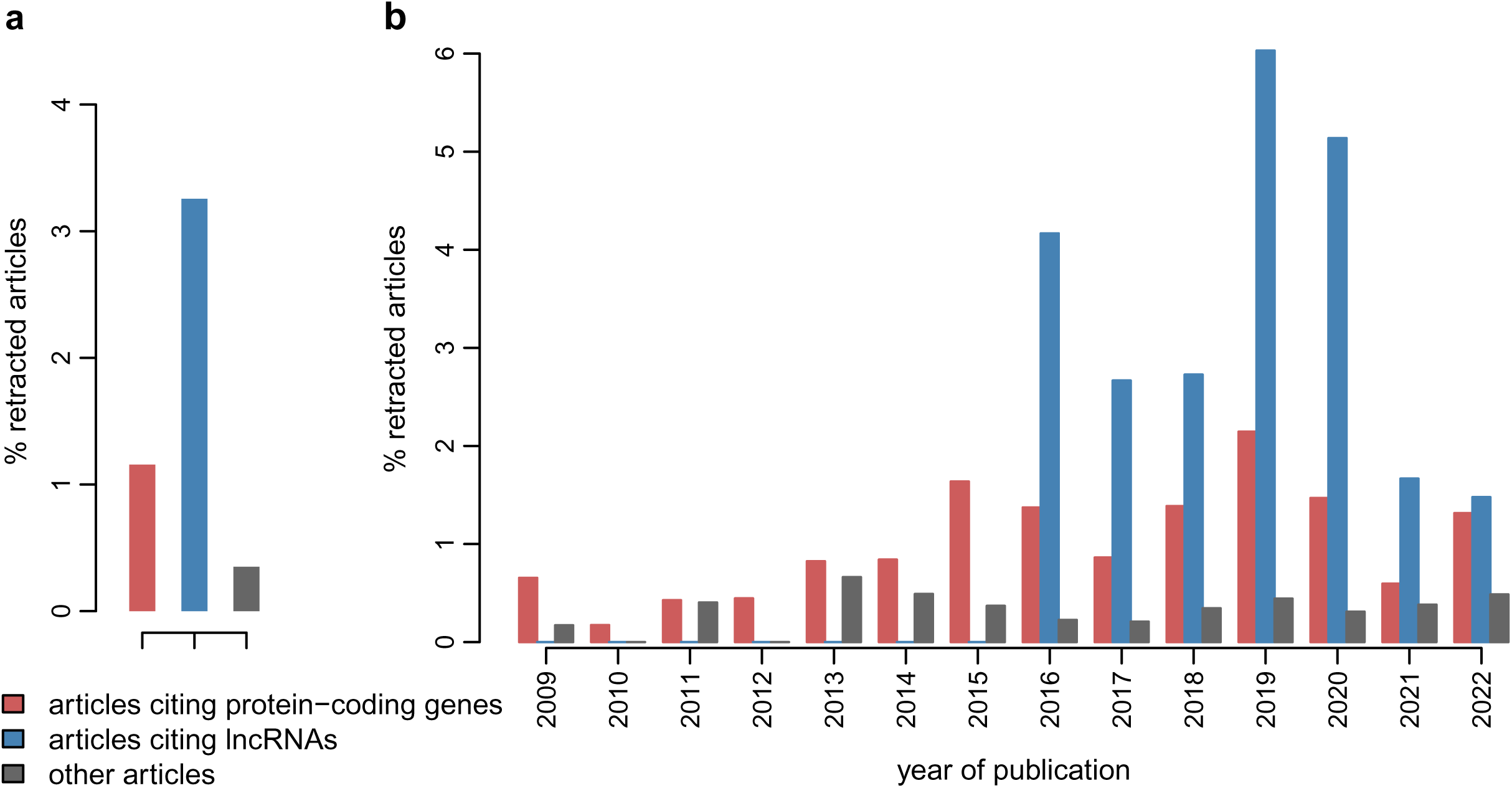
Retraction statistics for HCC-associated articles. a) Percentage of retracted articles, among all HCC-associated articles that cite protein-coding genes (red) or lncRNAs (blue) in the abstract, and among all other HCC-associated articles (gray). b) Percentage of retracted articles *per* year, between 2009 and 2022, for the same 3 article categories.

We also showed that the genomic and expression characteristics of HCC-associated lncRNAs are different from those of other lncRNAs. In particular, we found that HCC-associated lncRNAs are more highly expressed, more often found on the antisense strand of other genes, and more often transcribed from bidirectional promoters than other lncRNAs. These characteristics may not be independent from each other. The higher expression levels of HCC-associated lncRNAs can be easily explained, given that many of the studies that put them forward rely on comparative transcriptomics analyses, in particular differential expression analyses, the sensitivity of which is strongly correlated with gene expression levels (Love, Huber, and Anders 2014). Moreover, lncRNAs that are found on the antisense of protein-coding genes or transcribed from bidirectional promoters shared with protein-coding genes are generally more highly expressed than purely intergenic lncRNAs. Thus, the skewed genomic characteristics of HCC-associated lncRNAs are expected to some extent, given the discovery bias in favor of highly expressed genes. However, we also observed that HCC-associated lncRNAs have neighbors that are themselves frequently differentially expressed between tumors and controls, significantly more so than protein-coding genes. This could be in part due to the presence of large-scale copy number alterations in HCC, which could encompass several neighboring genes. This observation might also indicate that neighbors of genes that are already known to be associated with HCC may be more frequently investigated or more frequently put forward in publications, which would introduce a bias in the genomic landscape of lncRNAs associated with HCC. Evidently, and especially given that antisense lncRNAs are often proposed to regulate the expression of the gene they overlap with (Werner et al. 2024), pairs of sense/antisense genes can be genuinely involved in similar biological processes, including tumorigenesis. Nevertheless, we stress that the genomic context of the lncRNAs associated with HCC is not sufficiently discussed in the corresponding publications. For example, *HOTAIR* and *HOTTIP*, two of the 25 most frequently cited lncRNAs, are transcribed from the antisense strand of the HOXC and HOXA clusters of genes. However, we found that the publications that investigate the putative roles of *HOTAIR* and *HOTTIP* in HCC do not often investigate their protein-coding neighbors, despite their well described roles in organismal development (Supplementary Table 11). Some exceptions exist (Quagliata et al. 2014), but we believe that the genomic context of candidate disease-associated lncRNAs should systematically be assessed, in order to better evaluate their putative biological roles.

Finally, in this manuscript we analyzed in depth the previously reported expression patterns and biological roles for the 25 lncRNAs that were most frequently associated with HCC in scientific publications. These top lncRNAs include transcripts that have been discovered and extensively studied outside the HCC context, such as *H19*, *XIST*, *MALAT1*, *HOTAIR* (Figure 4). This may again be indicative of a biased representation of HCC-associated lncRNAs, as genes that are already well known tend to be more often investigated or put forward as interesting candidates. Strikingly, for 10 out of these 25 lncRNAs we found contradictions either within the existing literature, or between previous reports and our own findings regarding their expression patterns in HCC tumors and controls (Table 1). This is the case for some of the best-described lncRNAs, such as *MALAT1*, *NEAT1*, *HOTAIR*, *H19* and *XIST*. Some of the contradictions between the expression patterns observed for lncRNAs in different publications might be due to differences in the genetic background of the patient cohort, or in the techniques that were used to collect biological samples or to prepare RNA sequencing libraries. As lncRNAs are often proposed as candidate oncogenes or tumor suppressors starting from transcriptome comparative analyses, we believe it is crucial to analyze multiple patient cohorts, to better assess the impact of cohort composition (not only in terms of genetic background but also in terms of underlying diseases, tumor differentiation grades, etc.) on the observed expression patterns. Here, we endeavored to do this by analyzing two large patient cohorts, including the TCGA dataset that is often used as a reference in transcriptome-based studies. Importantly, the analyses that we performed on the two datasets are in agreement with each other and in contradiction with the literature for some of the most highly-cited lncRNAs, including *HOTAIR*, *H19* and *XIST* (Figure 4, Supplementary Figure 4). This comparison allows us to be confident that the inconsistencies that we observe between our observations and previous reports are not all simply due to differences among cohorts.

These inconsistencies that we observed for the expression patterns of lncRNAs do not necessarily invalidate the proposed biological roles of the lncRNAs, if - and only if - these roles are supported by additional, solid evidence. However, contradictions regarding the functions of lncRNAs were also previously reported following experimental validations of lncRNAs, as illustrated by the debates surrounding *MALAT1* (Kim et al. 2018) and *H19* (Tietze and Kessler 2020). More worryingly, our scan of the literature for the top 25 HCC-associated lncRNAs leads us to believe that differential expression between HCC tumors and controls is often the only basis for proposing oncogene or tumor suppressor roles for lncRNAs (Table 1, Supplementary Table 11). We also find it puzzling and concerning that the overwhelming majority (24 of 25) of these highly-cited lncRNAs are proposed to act as competing endogenous RNAs (Table 1), including lncRNAs such as *XIST* for which other biological mechanisms are well established (Augui, Nora, and Heard 2011).

A recent review has aimed to raise awareness of questionable evidence and unsubstantiated claims related to lncRNAs in the scientific literature (Ponting and Haerty 2022). We believe that this critical point of view is extremely important in the context of cancer research, which cannot afford to be plagued by unreliable reports of lncRNA functions.

## Materials and methods

### Gene annotations

We extracted human gene annotations from Ensembl release 109 (Martin et al. 2023). We selected long non-coding RNAs and protein-coding genes based on Ensembl gene biotypes (“lncRNA” and “protein_coding”, respectively). We excluded from our analyses genes that were located on alternate sequences (e.g., haplotypes corresponding to the MHC region).

### Literature analysis

We used the *esearch* tool from the NCBI E-utilities to search in PubMed for articles containing the phrase “hepatocellular carcinoma” in the title (hereafter named HCC-associated articles). We obtained 76,491 PubMed entries. The search was performed on March 11th, 2024. We processed the results with a custom Perl script to extract for each entry its PubMed identifier, the journal, the year of publication, the publication type, as well as the genes that were cited in the abstract. To identify the cited genes, we first extracted from Ensembl release 109 (Martin et al. 2023) HGNC symbols and synonyms for each gene. We then matched abstract words (after removing punctuation, parentheses, and brackets) with gene symbols and synonyms. We excluded 92 gene names that coincided with commonly used abbreviations not related to genes (e.g. HR for hazard ratio, CP for Child-Pugh cirrhosis score, etc.). For the 25 most-cited lncRNAs, we extracted all articles that cite them and manually extracted from the abstracts the following information, if available: the expression pattern in HCC tissues versus controls; the expression pattern in HCC model cell lines versus control cell lines; whether the article was focused on the lncRNAs (as opposed e.g. to articles focused on a different type of gene or on a treatment); whether the article was a review; whether the lncRNA was presented as an oncogene, a tumor suppressor or a biomarker; whether the lncRNA was presented as a competing endogenous RNA for microRNAs; whether the lncRNA was said to regulate microRNA expression. We then summarized this information for each lncRNA, excluding review articles and articles that were not focused on lncRNAs.

### Transcriptome sequencing data

We analyzed transcriptome sequencing data from two HCC patient cohorts. The first one, which is presented in the main analyses in the manuscript, was described in a recent publication (Ng et al. 2022). This dataset included transcriptome sequencing data for tumor biopsies from 114 HCC patients, obtained prior to treatment. The sequencing data is available at the European Genome-Phenome Archive with the accession number EGAS00001005074. We complemented this dataset with transcriptome sequencing data for adjacent tissue biopsies for the same cohort of patients. The sequencing data was generated with the same methods as the corresponding tumor samples (Ng et al. 2022). Briefly, RNA was extracted with the RNA MiniPrep Plus kit (Zymo Research), RNA-seq libraries were prepared with the Illumina TruSeq Stranded Total RNA Library Prep Kit with Ribo-Zero Gold and sequenced as single-end 126 base pairs (bp) reads on an Illumina HiSeq 2500 machine. We were able to analyze transcriptome sequencing data for 105 patients for which paired tumor and adjacent tissue biopsies were available. We submitted the transcriptome sequencing data for the adjacent tissue biopsies to the European Genome-Phenome Archive (accession number pending). The second HCC patient cohort was generated by The Cancer Genome Atlas (TCGA) consortium (The Cancer Genome Atlas Research Network 2017). Specifically, we analyzed TCGA transcriptome sequencing data for paired tumor and adjacent tissue biopsies, for 37 HCC patients. This transcriptome sequencing dataset was generated with the Illumina TruSeq mRNA protocol, which selects polyA-containing transcripts and does not preserve RNA strand information.

### Gene expression estimation

We used *kallisto* version 0.46.1 (Bray et al. 2016) to compute expression levels for Ensembl-annotated genes. For the Ng et al 2022 single-end RNA-seq data, we set the average RNA fragment length at 200 bp and the standard deviation at 25 bp and we set the strand-specific library orientation parameter to “rf-stranded”. For the TCGA data, which consists of paired-end unstranded reads, these settings were not used. For both Ng et al 2022 and TCGA data we enabled the bias correction method implemented in *kallisto* to account for the effects of nucleotide composition on the frequency and distribution of RNA-seq reads, using a patch to correct an error in the bootstrap procedure when bias correction is enabled (https://github.com/pveber/kallisto/tree/use-bias-in-bootstrap).

### Differential expression analyses

We used the *tximport* R package (Soneson, Love, and Robinson 2015) to import in R the gene expression estimates obtained with *kallisto*. We then used functions in the DESeq2 R package (Love, Huber, and Anders 2014) to test for differential expression (DE) between conditions. For both Ng et al 2022 and TCGA datasets, we tested for DE between tumor and adjacent tissue samples. To do this, we constructed an additive model in DESeq2 with the tissue type (tumor or adjacent tissue) and the patient identity as explanatory variables and we used the Wald test to evaluate expression differences between tissue types. For the Ng et al 2022 dataset, we also tested for DE between tumor differentiation stages, by contrasting tumor samples with Edmondson-Steiner grades I and II and tumor samples with Edmondson-Steiner grades III and IV. For this analysis, we again constructed an additive model with two explanatory variables, namely the sex of the patient and the Edmondson-Steiner grade class, and we applied the Wald test to evaluate DE between Edmondson-Steiner grade classes. For all DE analyses, we used the Benjamini-Hochberg procedure to correct for multiple testing and we selected significantly DE genes by setting a maximum false discovery rate (FDR) threshold.

### Principal component analyses

We used the *dudi.pca* function in the *ade4* R package to perform principal component analyses (PCA). We used as an input TPM values, transformed with the function x −> log2(x+1). The data was centered before performing the PCA. We computed the percentage of variation explained by each PCA axis starting from the eigenvalues corresponding to each axis.

### Gene localisation analysis

We extracted gene coordinates from Ensembl release 109 (Martin et al. 2023) and we analyzed them to extract gene body overlaps on the sense and antisense strand. We then looked for bidirectional promoters by extracting the positions of the transcription initiation site (TSS) for each gene and identifying the TSS found on the opposite strand, within a maximum distance of 1 kilobase (kb).

### Sequence conservation analyses

We downloaded PhastCons (Siepel et al. 2005) sequence conservation scores from the UCSC Genome Browser (Raney et al. 2024). These scores were computed on a whole genome alignment between human (hg38 assembly) and 29 other mammals, including 26 primates. We computed the average PhastCons score for the exonic regions of each gene, excluding exonic parts that overlap with exons from other genes.

### Ethics

Human tissues were obtained from patients undergoing diagnostic liver biopsy at the University Hospital Basel between 2008 and 2018. Written informed consent was obtained from all patients. The study was approved by the ethics committee of the northwestern part of Switzerland (Protocol Number EKNZ 2014-099). All research was conducted in accordance with both the Declarations of Helsinki and Istanbul.

## Supplementary Figure legends

**Supplementary Figure 1.**
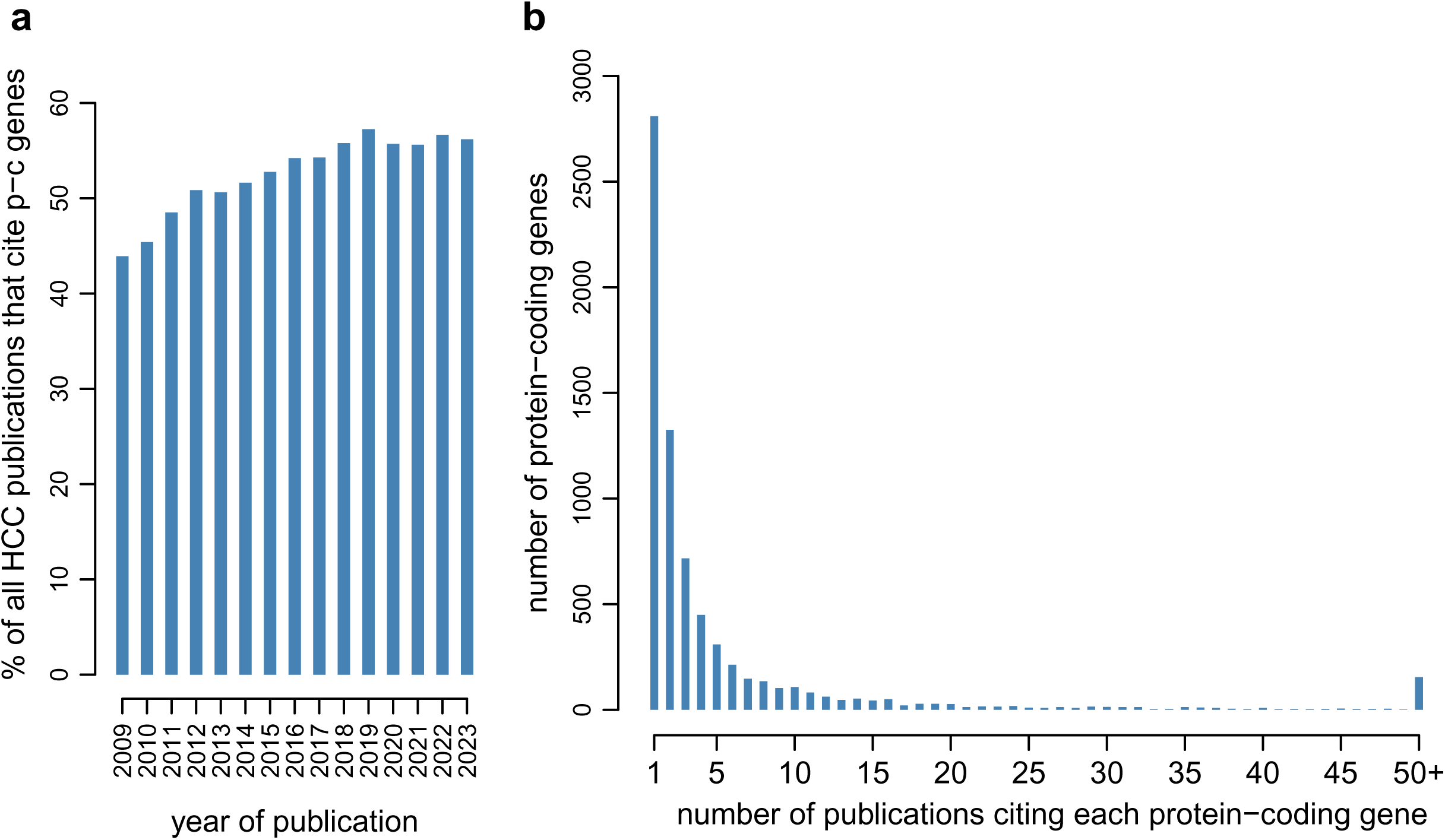
HCC-associated publications that cite protein-coding genes in the abstract. a) Barplot representing the percentage of all HCC publications that cite protein-coding genes in the abstract, between 2009 and 2022. b) Histogram of the number of HCC-associated publications in which each protein-coding gene is cited in the abstract.

**Supplementary Figure 2.**
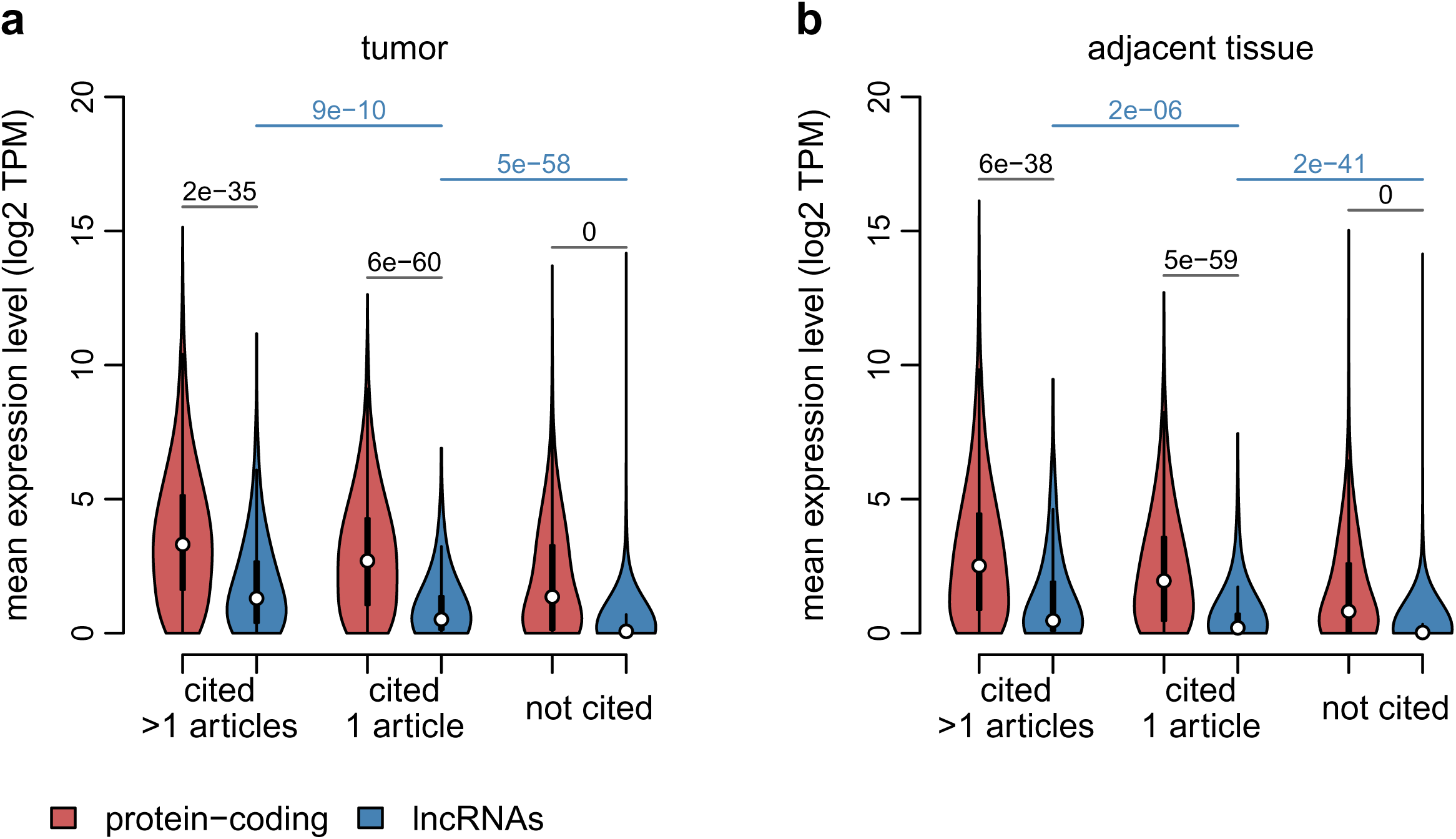
Expression patterns for HCC-associated lncRNAs, in comparison with protein-coding genes and other lncRNAs, in the TCGA HCC dataset. Protein-coding genes are depicted in red and divided into 3 categories: cited in more than one HCC publication, in exactly one publication, or not cited. Long non-coding RNAs are depicted in blue and divided in the same categories. Red: protein-coding genes. Blue: lncRNAs. a) Violin plots representing the distribution of the average expression level (log2-transformed TPM) for TCGA tumor samples. b) Same as a), for TCGA adjacent tissue samples. Numeric values represent p-values of the Wilcoxon rank sum test, between protein-coding genes and lncRNAs or between citation classes for lncRNAs.

**Supplementary Figure 3.**
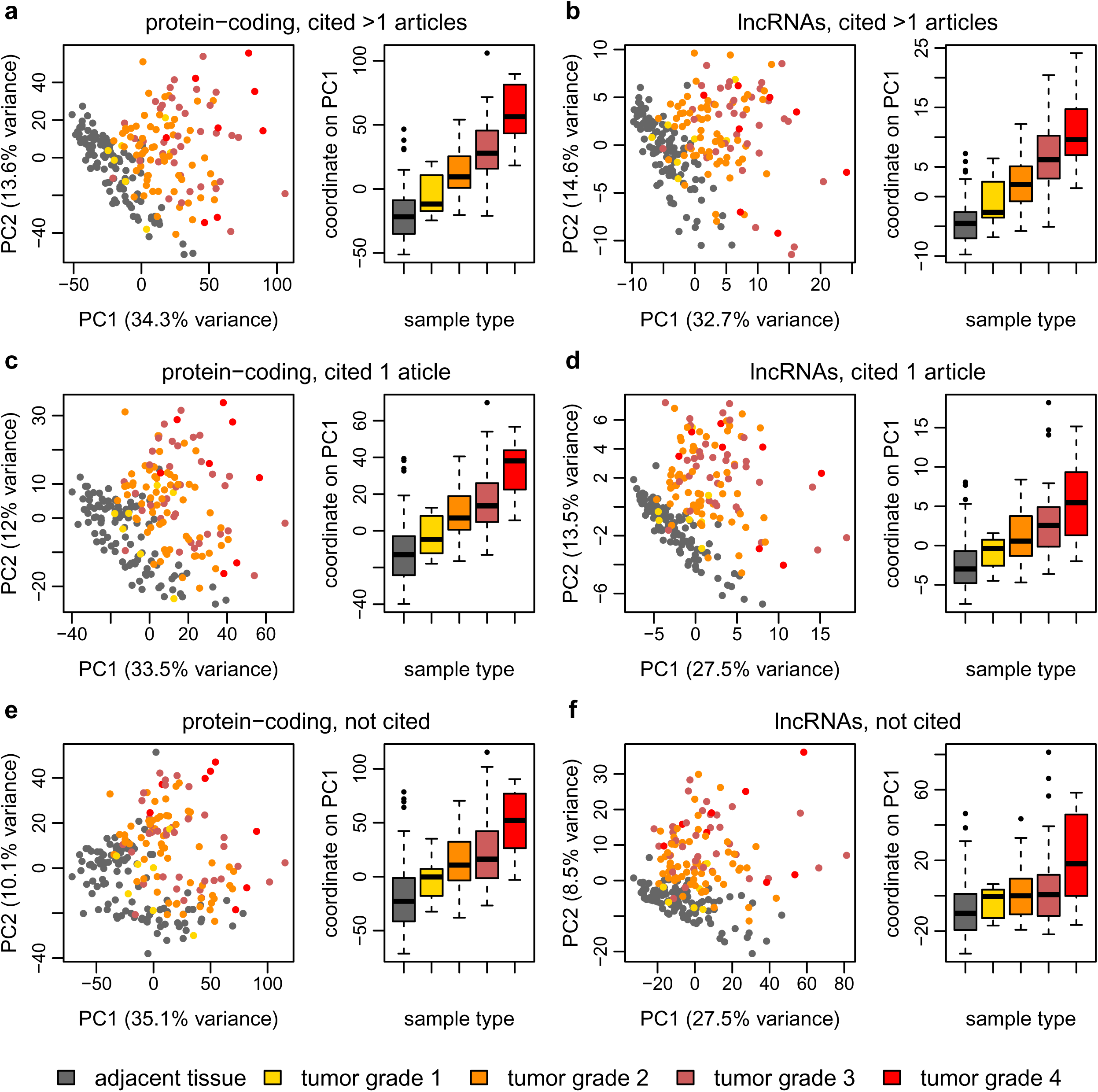
Principal component analyses (PCA) for lncRNA and protein-coding genes expression patterns, for the Ng et al 2022 transcriptome dataset. PCA was performed separately for each class of genes, on log2-transformed TPM values. Dots represent individual samples. Gray: adjacent tissue samples; yellow: tumor samples with Edmondson-Steiner grade I; orange: tumor samples with Edmondson-Steiner grade II; maroon: tumor samples with Edmondson-Steiner grade III; bright red: tumor samples with Edmondson-Steiner grade IV. a) PCA first factorial map, for protein-coding genes cited in more than one HCC publication. b) Same as a), for lncRNAs cited in more than one HCC publication. c) Same as a), for protein-coding genes cited in exactly one HCC publication. d) Same as a), for lncRNAs cited in exactly one HCC publication. e) Same as a), for protein-coding genes not cited in association with HCC. f) Same as a), for lncRNAs not cited in association with HCC.

**Supplementary Figure 4.**
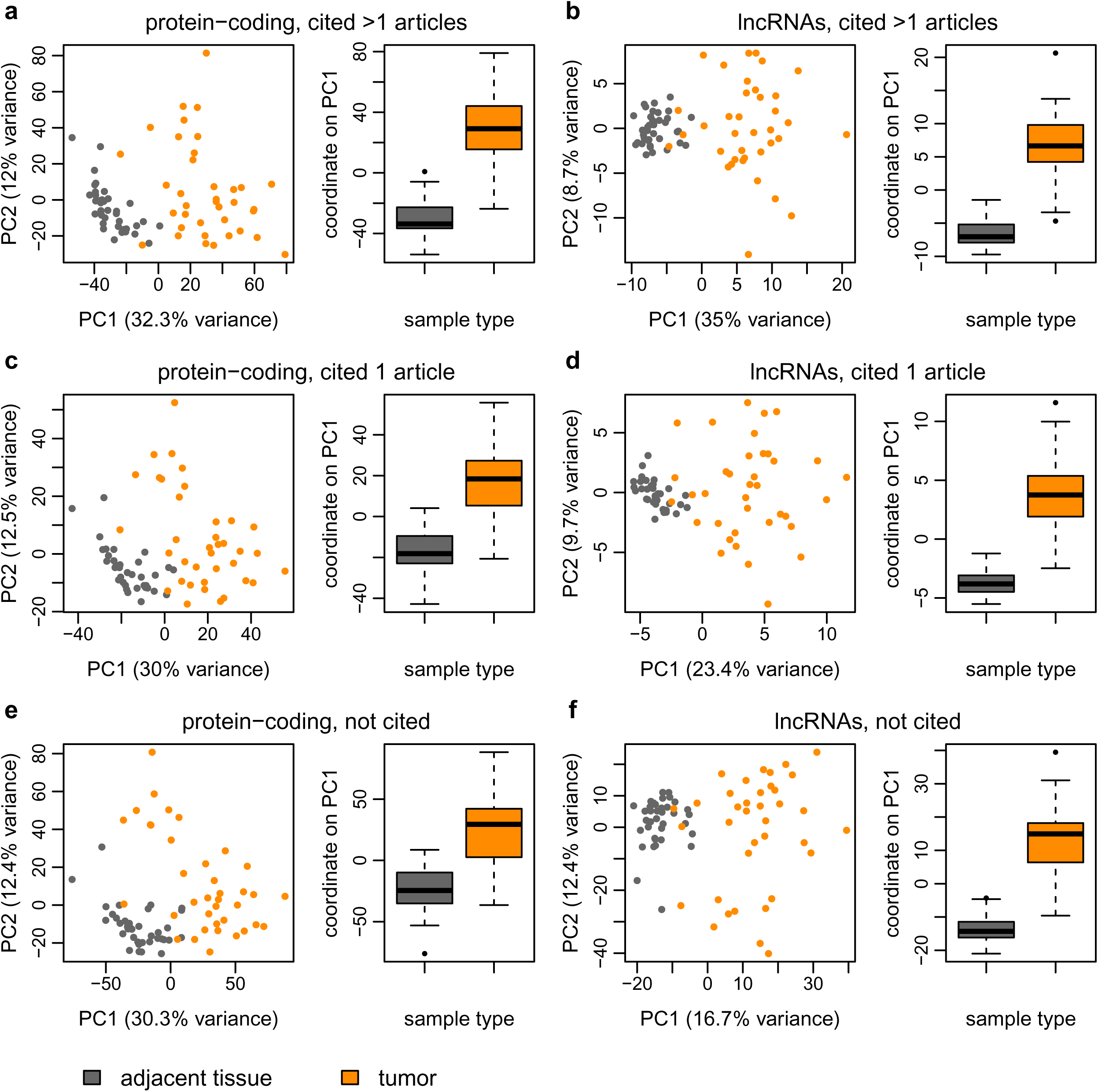
Principal component analyses (PCA) for lncRNA and protein-coding genes expression patterns, for the TCGA transcriptome dataset. PCA was performed separately for each class of genes, on log2-transformed TPM values. Dots represent individual samples. Gray: adjacent tissue samples; orange: tumor samples. a) PCA first factorial map, for protein-coding genes cited in more than one HCC publication. b) Same as a), for lncRNAs cited in more than one HCC publication. c) Same as a), for protein-coding genes cited in exactly one HCC publication. d) Same as a), for lncRNAs cited in exactly one HCC publication. e) Same as a), for protein-coding genes not cited in association with HCC. f) Same as a), for lncRNAs not cited in association with HCC.

**Supplementary Figure 5.**
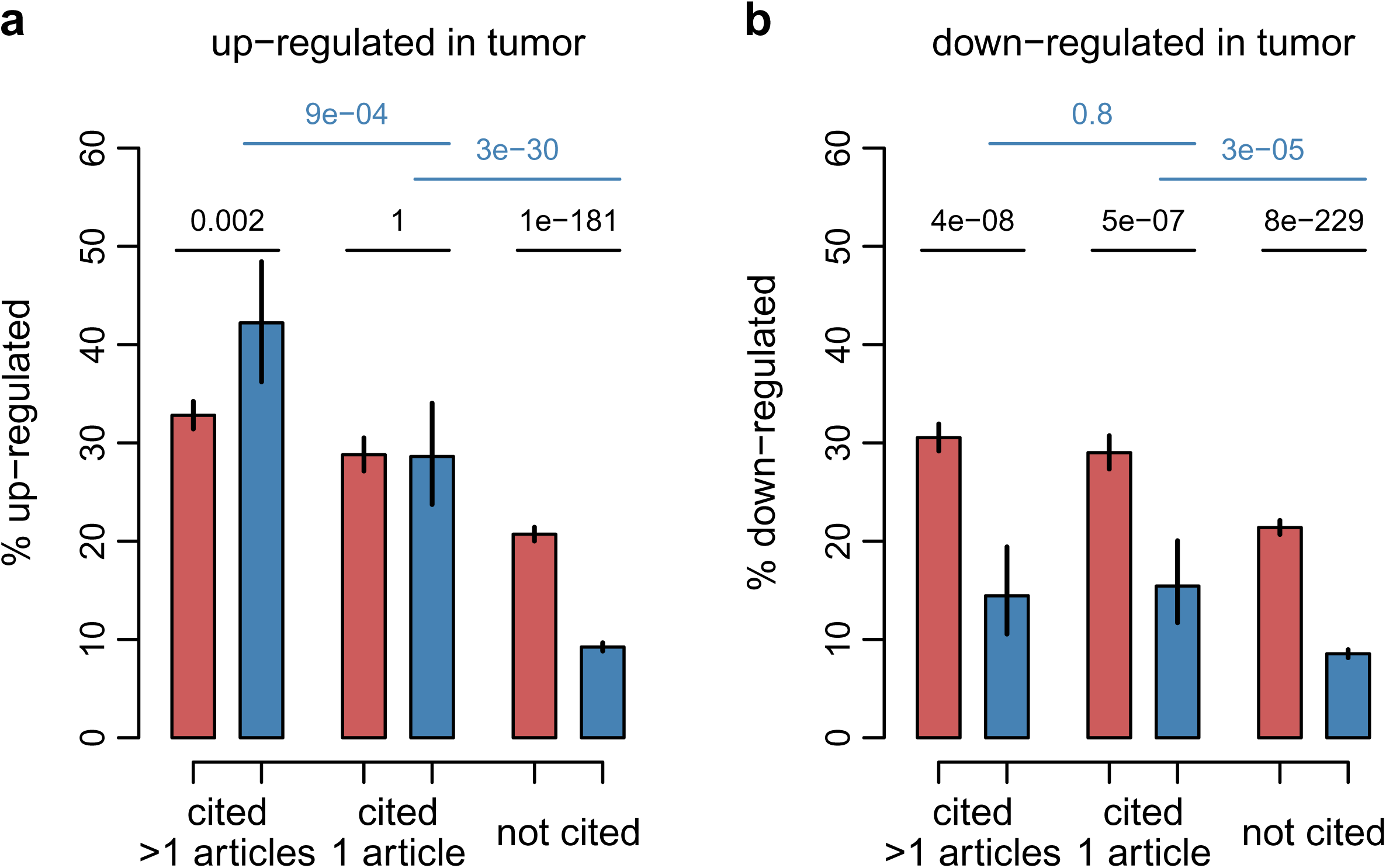
Differential expression patterns for HCC-associated lncRNAs and comparison with other lncRNAs and with protein-coding genes, in the TCGA transcriptome dataset. Protein-coding genes are depicted in red and divided into 3 categories: cited in more than one HCC publication, in exactly one publication, or not cited. Long non-coding RNAs are depicted in blue and divided in the same categories. A maximum FDR threshold was set to extract significantly DE genes. a) Percentage of genes that are significantly up-regulated in tumors compared to adjacent tissues. b) Percentage of genes that are significantly down-regulated in tumors compared to adjacent tissues. Vertical bars represent 95% confidence intervals.

**Supplementary Figure 6.**
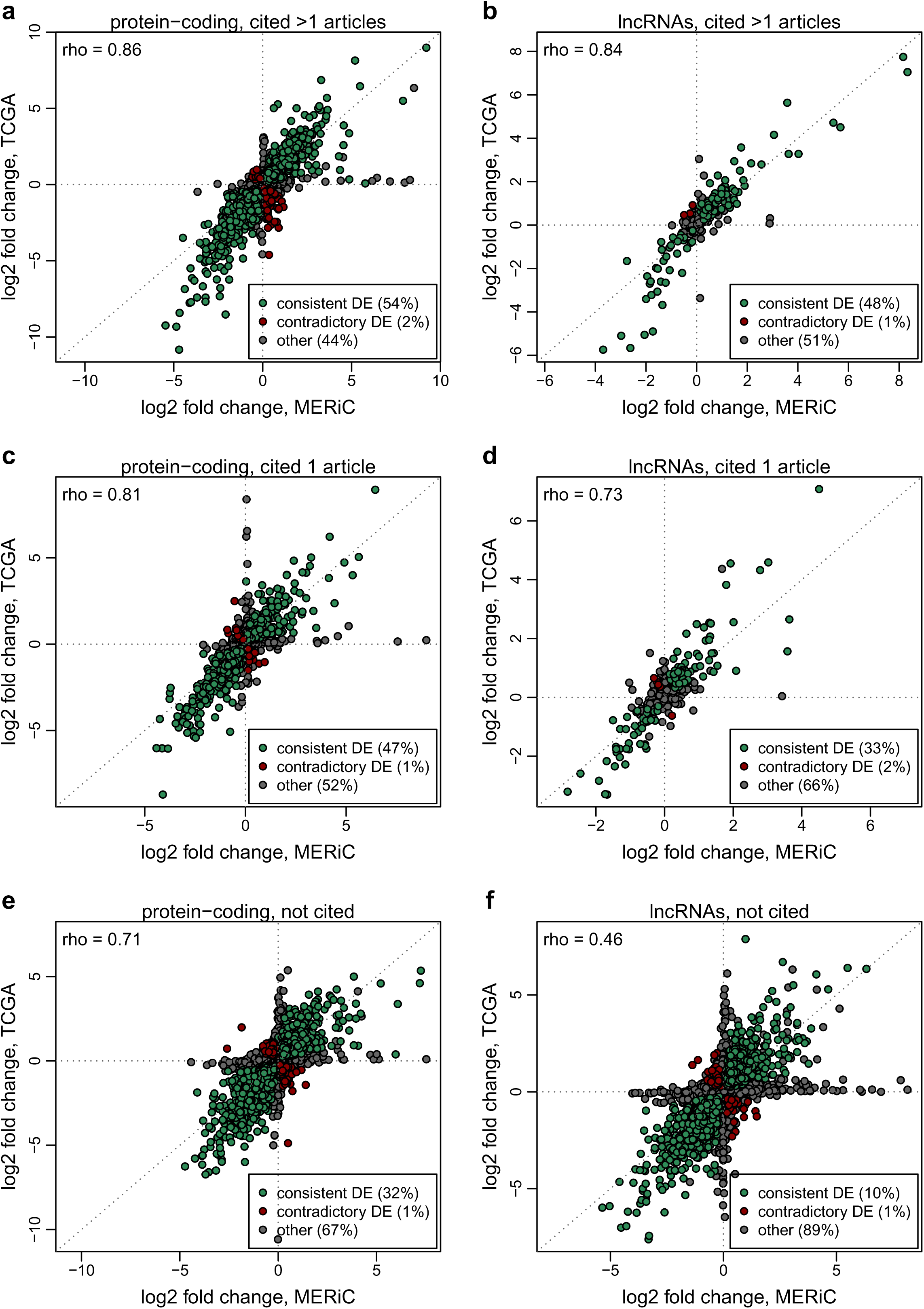
Comparison of DE patterns between the Ng et al 2022 and the TCGA datasets. Protein-coding genes and lncRNAs are divided into 3 categories: cited in more than one HCC publication, in exactly one publication, or not cited. Green dots represent genes with consistent DE patterns (FDR < 5% in both datasets, in the same direction). Red dots represent genes with contradictory DE patterns (FDR < 5% in both dataset, opposite directions). Gray dots represent all other cases. a) Scatter plot representing the relationship between the log2 fold expression change between tumors and adjacent tissues in the Ng et al dataset (X-axis) and in the TCGA dataset (Y-axis), for protein-coding genes cited in more than one HCC-associated publication. b) Same as a), for lncRNAs cited in more than one HCC-associated publication. c) Same as a), for protein-coding genes cited in exactly one HCC-associated publication. d) Same as a), for lncRNAs cited in exactly one HCC-associated publication. d) Same as a), for protein-coding genes not cited in association with HCC. e) Same as a), for protein-coding genes not cited in association with HCC.

**Supplementary Figure 7.**
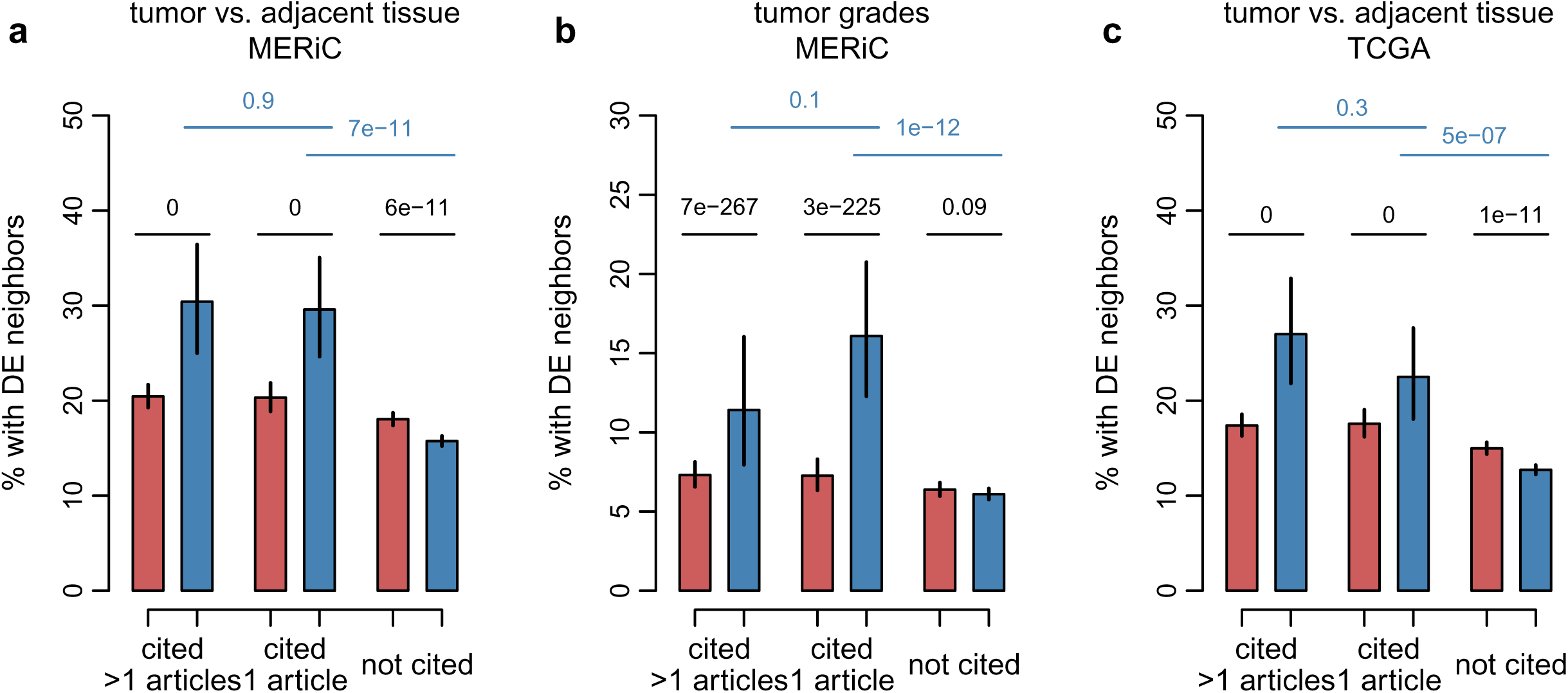
Neighbors of HCC-associated lncRNAs are often differentially expressed. Protein-coding genes are depicted in red and divided into 3 categories: cited in more than one HCC publication, in exactly one publication, or not cited. Long non-coding RNAs are depicted in blue and divided in the same categories. a) Percentage of genes that have a bidirectional promoter neighbor that is significantly DE between tumors and adjacent tissues in the Ng et al 2022 dataset. b) Percentage of genes that have a bidirectional promoter neighbor that is significantly DE between tumors with Edmondson-Steiner grades III and IV and tumors with Edmondson-Steiner grades I and II in the Ng et al 2022 dataset. c) Percentage of genes that have a bidirectional promoter neighbor that is significantly DE between tumors and adjacent tissues in the TCGA dataset. Numeric values represent p-values of the Chi-squared test for the comparisons between protein-coding genes and lncRNAs, or between citation classes for lncRNAs.

**Supplementary Figure 8.**
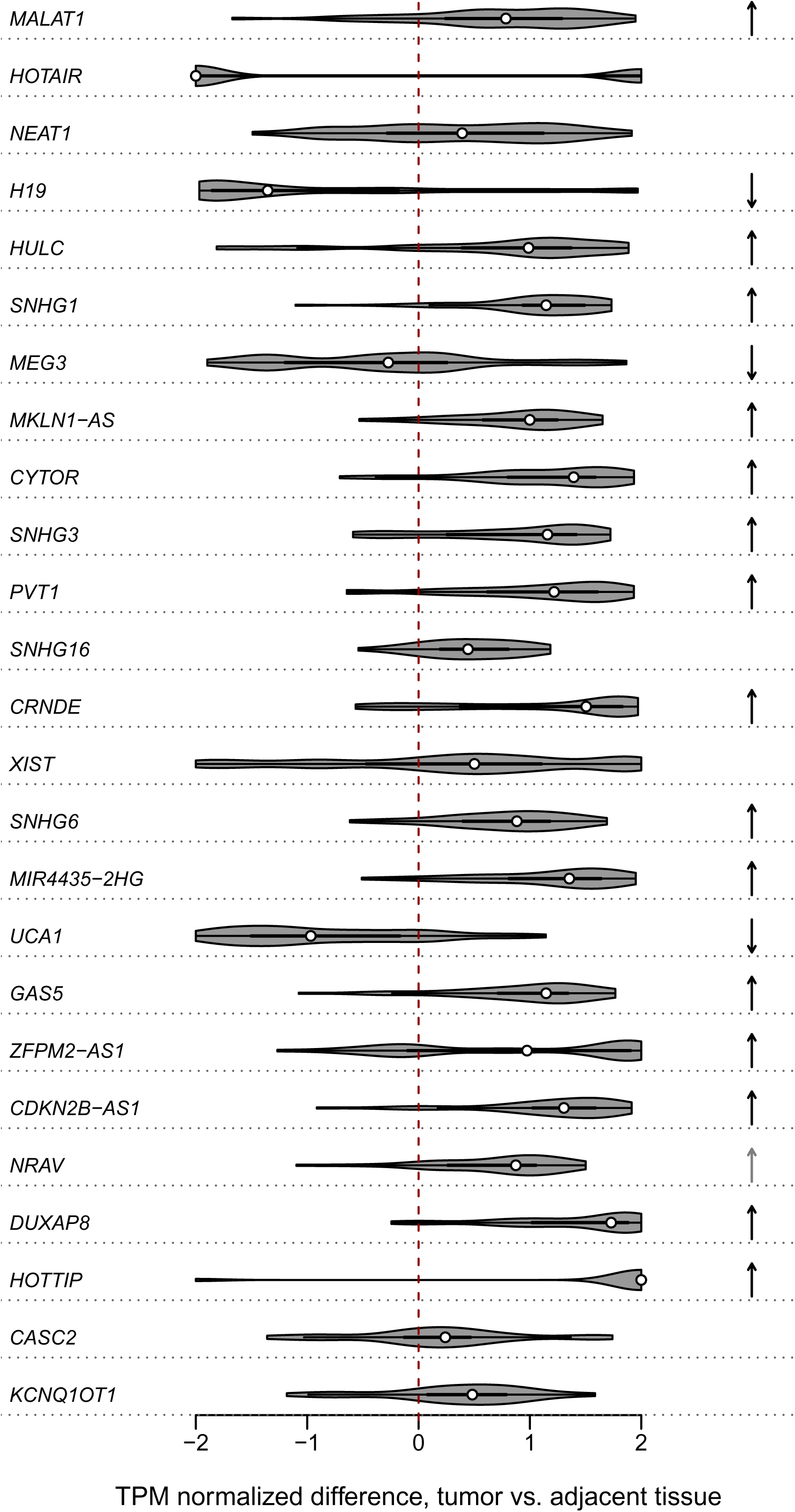
Expression patterns for the 25 lncRNAs with the highest numbers of HCC-associated citations in the TCGA transcriptome dataset. a) Violin plot depicting the distribution of the normalized expression differences between paired tumor and adjacent tissue biopsies, across the 105 analyzed patients. b) Arrows indicating whether the difference between tumor and paired adjacent tissue biopsies was statistically significant (FDR < 5%), and if so, the direction of the expression change (up-regulated or down-regulated). Dark gray arrows indicate absolute fold expression changes above 1.5.

## Supplementary Table legends

**Supplementary Table 1**. List of all PubMed articles that were analyzed here.

**Supplementary Table 2**. List of all HCC-associated lncRNAs.

**Supplementary Table 3**. List of all HCC-associated protein-coding genes.

**Supplementary Table 4**. Information for tumor biopsies derived from (Ng et al. 2022).

**Supplementary Table 5.** Information for adjacent tissue biopsies, paired with tumor biopsies derived from (Ng et al. 2022).

**Supplementary Table 6**. Genomic and expression characteristics of lncRNAs.

**Supplementary Table 7**. Genomic and expression characteristics of protein-coding genes.

**Supplementary Table 8**. Differential expression results, tumor vs. adjacent tissue biopsies, data from (Ng et al. 2022).

**Supplementary Table 9.** Differential expression results, Edmondson-Steiner grades, data from (Ng et al. 2022).

**Supplementary Table 10**. Differential expression results, tumor vs. adjacent tissue biopsies, data from TCGA.

**Supplementary Table 11**. Literature scan for the top 25 HCC-associated lncRNAs.

